# Impact of three light smoltification regimes on performance and genetic parameters of traits in Atlantic salmon

**DOI:** 10.1101/2024.04.02.587721

**Authors:** Bjarne Gjerde, Solomon Antwi Boison, David Hazlerigg, Trine Ytrestøyl, Turid Mørkøre, Even Jørgensen, Anja Striberny, Simen Rød Sandve

## Abstract

2800 Atlantic salmon pre-smolts (50 g on average, the offspring of 53 sires and 100 dams) were individually tagged with PIT-tags and distributed among six circular 1200L tanks with approximately 450 fish per tank. Fish in duplicated tanks were put on three different light regimes, i.e. six weeks on either 8L:16D, 12L:12D or 24L:0D followed by six weeks on 24L:0D. One week prior to their transfer as 1+ smolt to a net cage in the sea in June 2021 their body weight, length, subjectively scored smolt status and fin damage were recorded. Recording of animal traits (body weight, wounds, fin damage, snout damage) were also performed in November 2021 and in April 2022. During the light regimes, fish on the two short day regimes had slower growth compared to fish on continuous light. However, after four (November 2021) and ten (April 2022) months in the sea the effect of light regime on fish size was not significantly different from zero (P>0.05). The fish on the 24L:0D regime showed increased mortality from day two after sea transfer with an accumulated recorded mortality of 8.9% during the two first months while it was only 1.0% and 0.7% for the 12L:12D and the 8L:16D fish, respectively. However, from the third until ten months in the sea recorded mortality was very similar for fish on the three light regimes. The effect of light regime on the recorded welfare traits (fin and snout damage and runts) was not significantly different from zero. For traits measured prior to seawater transfer the difference between fish that survived and those that died during the first two months in the sea were largest in the 24L:0D group indicating a positive effect of the short-day regimes also on the general smolt synchronization (i.e., group level uniformity). Moderate heritability estimates were found for the external smolt indicator traits condition factor, smolt status score and skin silveriness, as well as for snout damage, wounds, runts and body weight, but low estimates fin damage. For survival in the sea heritability on the liability scale was 0.09 after four months in the sea and 0.22 from five to ten months in the sea. The estimated genetic correlations between the same trait of the three different light regimes were moderate to high and thus unimportant genotype by light regime interaction. The genetic correlation of body in June 2021 with survival after two months in the sea was high (0.96), but not significantly different from zero with survival after four months in the sea and survival from five to ten months in the sea. Genetic correlations of survival with the other traits recorded in June 2021, November 2021 and April 2022 were low to medium in magnitude and not significantly different from zero. Therefore, for genetic improvement of survival in the seawater period direct selection for increased survival and growth during the first months is probably a better strategy than to perform indirect selection for smolt indicator traits.

## 1. Introduction

Atlantic salmon (*Salmo salar*) is the major aquaculture commodity in Norway. In 2022 the production was 1.552 million tons (94% of the total Norway’s aquaculture production) valued to 100.8 billion NOKs (https://www.fiskeridir.no). Wild salmon is an anadromous fish adapted to spend their first years in fresh water, migrate to sea as ‘smolts’, grow to reach sexual maturity and finally return to the river of birth to spawn. Hence, the production of farmed salmon involves two phases; first a freshwater phase from the time of egg fertilization up to the smolt stage (12 to 18 months), when they are moved to floating seawater net cages where they grow to slaughter weight (1-1.5 years). To support the large salmon industry, >300 million smolts are reared and transferred to sea cages in Norway every year (304 mill in 2021). However, as many as 16.1% - a staggering 50 million fish - die before they reach the desired slaughter weight (Sommerset et al., 2022), and a significant part of this mortality takes place during first months after seawater transfer and is attributed to sub-optimal smolt development and physiology (Aunsmo et al., 2008). Mortality obviously represents economic loss for the farmers due to both reduced revenue and higher cost per kg fish slaughtered mainly due to reduced feed efficiency and thus high feed cost, but also a huge animal welfare problem. A pressing challenge in salmon aquaculture research is therefore to better understand smolt production factors that reduce mortality, increase feed resource efficiency and animal welfare post seawater transfer (Kolarevic et al., 2014; Ytrestøl et al., 2020; Khaw et al., 2021; Striberny er al., 2021).

The developmental process turning a freshwater salmon (parr) to a saltwater adapted fish (smolt) is termed ‘smoltification’. This encompasses a multitude of behavioural, morphological, and physiological preparatory changes that helps the fish cope with the seawater environment. Skin colour changes from a darker tone with parr marks (i.e. spots) to a silvery appearance and the body shape becomes longer and thinner, resulting in a reduction in condition factor (Folmar et al., 1998; Hoar et al., 1988; Sigholt et al., 1998; McCormic et al., 2000; Nichols et al., 2008; Piironen et al., 2013; Khaw et al., 2021). At the molecular level many organs change functional properties (refs to liver, kidney, gut papers), and many of these changes are related to the change from a freshwater to a seawater rearing environment. In full strength seawater, the osmolarity of the fish internal fluids is around one-third of the surrounding water (Evans et al., 2005), hence they become hypoosmotic relative to the external seawater (McCormick et al., 2013). To cope with this, hypo-osmoregulatory ability is developed during smoltification through changes in kidney function (master thesis), enhanced water permeability in the intestine (Boeuf, 1993; Ura et al., 1997; Duarte et al, 2023), and remodelling of gill cell populations (McCormick et al., 2013), resulting in more efficient salt excretion capacity.

In nature smoltification is triggered by increased daylength in the spring, making the salmon prepared to migrate to sea in early summer (May-June in Northern Hemisphere). Production of smolts in aquaculture has therefore traditionally been achieved by mimicking a natural photoperiodic regime. Typically, these production protocols use continuous light in the parr (pre-smolt) stage, interrupted by 6-8 weeks with short days (≤12 h of daylight), followed by 6-8 weeks on 24 h daylight. However, motivated by growth maximization (Bjornsson et al., 1989; Myklatun et al., 2023) and practical considerations salmon farming industry has more recently explored alternative smolt production protocols. These protocols include rearing the smolts to a much large body weight (200-500 g as compared to 75-200 g) due to the positive correlation between size and salinity tolerance (Handeland et al., 2013; Sigholt et al., 1995; Strand et al., 2018), and the use of feeds with added salt prior to seawater transfer which is known to increase seawater tolerance (Basulto, 1976; Salman and Eddy, 1988; Staurnes and Finstad, 2000; Duarte et al., 2023; Myklatun et al., 2023). Few recent trials have evaluated smolt performance under different rearing protocols.

Ytrestøyl et al. (2022) compared the growth of three groups of Atlantic salmon in RAS that were exposed to either a conventionally 12L:12D, 12L:12D or a 24L:0D light smoltification regime, respectively prior to transfer to net-cages in seawater at an average body weight of 100, 200 and 600 g, respectively. At first sampling eight weeks after seawater transfer the group exposed to the 24L:0D regime was bigger than the 200 g group exposed to 12L:12D regime because of the higher growth rate prior to sea transfer. However, the best performing group (largest average harvest weight, low mortality, and no sexual mature fish) was the 100 g group smoltified under the 12L:12D regime. Harvest weight was lowest in fish transferred to sea at 600 g, despite having the highest day-degree sum during their life span. These results strongly indicate that smolts reared under constant light experience difficulties to adjust to the new saline environment.

In another trial Striberny et al. (2021) compared the performance of smolts exposed to either a short-day (7L:14D) or raised under constant light (24L:0D), in combination with transition-feed. Here the authors confirmed that salt-spiked transition feed increases ability to hypo-osmoregulate, but that the combination of short-day treatment and transition-feed resulted in the best performing smolts (based on growth in seawater). Taken together; smolt production protocols matters and these protocols benefits from leveraging seasonal signals which salmon is naturally adapted to respond to.

In addition to environmental factors, genetic variation is also known to play a major role in quantitative trait variation. Nevertheless, genetics has mostly been neglected in research on smolt development and rearing protocols. Direct selection for improved smolt characteristics can only be obtained if the actual trait(s) show genetic variation, and their genetic correlation to other important production traits (e.g., growth and survival) should preferably be favorably, and not strongly unfavorable as this would result in less genetic gain for the other directly or indirectly selected for. Little is known about the magnitude of genetic variation in smolt development characteristics and their genetic and environmental correlation to other traits. In addition, no estimate is available about to what degree families rank differently with respect to important smolt traits as well as other production traits when sibs of the same families are smoltified on different light regimes, i.e., the degree of genotype by light regime interaction. In a rare and pioneering study on smolt genetics Khaw et al. (2021) found significant heritability of a molecular marker for smoltification status (the NKA SW/FW ratio), indicating that selection could achieve a more synchronized smoltification process. However, this study was limited in that it did not contrast production protocols with different day-lengths and did not track individual fish from the freshwater stage to their performance (e.g., growth, survival) in seawater.

The objectives of the present study were therefore to estimate the impact of three different light smoltification regimes on (a) growth, survival and animal welfare traits in the seawater phase of Atlantic salmon, (b) the magnitude of the genetic variation in the traits, and (c) on the magnitude of the genetic and environmental correlations among the traits.

## 2. Material and Methods

### 2.1. Ethical statement

The experiment was performed according to EU regulations concerning the protection of experimental animals (Directive 2010/63/EU). Appropriate measures were taken to minimize pain and discomfort. The experiment was approved by the Norwegian Food and Safety Authority (FOTS id. number 25658).

### 2.2. Freshwater phase

#### 2.2.1. Fish

The fish material originated from a total of 5,000 eyed-eggs of the Mowi strain from 100 full-sib families (50 eggs per family that were pooled at the eyed-egg stage). The families were produced over a three-week period in November 2019 but were temperature controlled to the same sum day-degree at the eyed-egg stage in January 2020. Of the 100 families 94 were the offspring of 47 sires and 94 dams produced using a nested 1 sire : 2 dam mating ratio, while the remaining six families were produced using a 1 sire : 1 dam mating ratio.

#### 2.2.2. Rearing conditions prior to treatment start

The 5000 eyed eggs were shipped from Mowi Genetics AS, Bergen to the Aquaculture Research Station in Tromsø (ARST) on 28.01.2020 and incubated in one 200 L freshwater tank at 1.8 °C and that was increased gradually to 7.9 °C within the first four days. Hatching started on 11.02.2020 and was completed after one week. The hatchlings were startfed on 01.04.2020 at which the water temperature was gradually increased to 10-12 °C and the light regime changed from 24 h darkness to 24 h light. From 19.05.2020 water temperature was changed to ambient temperature (4°C-12°C) until September 2020, when ambient temperature naturally decreased and kept constant at 8 °C.

Malformed juveniles were constantly removed and on 22.07.2020, 4090 fish were transferred to a 1200 L tank. On 15.12.2020, fish that were smaller than 10 g (“social loosers”) were sorted out and the remaining 3200 fish (mean body weight 31.4 g, SD=9.7 g) were redistributed among four 1200 L freshwater tanks. In early February the fish were vaccinated by Alpha Ject micro 6 (PHARMAQ®) against *Aeromonas salmonicida, Listonella anguillarum O1 and O2a, Vibrio salmonicida, Moritella viscosa* and *IPN virus Sp*.

The fish were fed *ad libitum* with commercial extruded pellet feed (Nutra Sprint, 0.5 mm and 1 mm and Nutra Olympic 1.5 mm, 2 mm, and 3 mm from Skretting AS, Stavanger, Norway) using automatic feeders. Pellet size was adjusted to match fish size according to the manufacturer protocol.

#### 2.2.3. Tagging and tissue sampling

In March 2021 (15 to 23) a total of 3000 fish (approximately 50 g) were individual tagged with passive integrated transponders (PIT-tag), fin-clipped and distributed among six circular 1200 L tanks with 500 fish per tank). Their body weights at tagging were not recorded. Prior to this the fish were fasted for 24 hours and anesthetized in benzocaine (60 ppm). The tags (ID-100A, www.trovan.com) were surgically implanted into the left intraperitoneal cavity using syringes pre-loaded with transponders, and a small incision made with a scalpel. Subsequently, a biopsy of the adipose fin was taken with scissors and stored in 96-tube-racks consisting of 1 ml barcode labelled tubes, containing 750 μl 100 % ethanol. Finally, the fish was released into a wake-up bath and transferred to one of six tanks. Biopsies were stored at room temperature until shipping for genotyping.

#### 2.2.4. Rearing conditions during smoltification

For a period of six weeks, from 22. March to 3. May 2021, the fish in duplicate tanks were subjected to three different photoperiodic light regimes (8L:16D, 12L:12D, 24L:0D). Water temperature was kept constant at 8 °C followed by 6 weeks on a 24L:0D light regime. The fish in all tanks were fed ad libitum with extruded salmon pellet feed (Nutra Olympic 3 mm, Skretting AS) for 8 hours of light the using automatic feeders.

#### 2.2.5. Smolt sampling

Recording of external smolt characters prior to seawater transfer was carried out from 7. To 14. June 2021. Fish were fasted 24h and the sampling of fish was randomized and carried out tank by tank. Fish were anesthetized with benzocaine (60 ppm) and the following traits recorded (see Table 1): Body weight (to the nearest g), body (fork) length (to the nearest 0.5 cm), fin damage (present/absent), and skin silveriness as a measure of smolt status (assessed visually according to the following criteria 1 = parr marks clearly visible, 2= silvery appearance but still visible parr marks, or 3= silvery skin, dark fin edges and no parr marks visible). All traits were recorded electronically using Fishreader and the software ZeusCapture (Trovan Ltd.). The proportion of fish with silvery skin was calculated as the ratio of the number of fish with score 3 and the total number of fish with score 1, 2 and 3. Condition factor was calculated as: 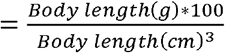 .

**Table 1.**
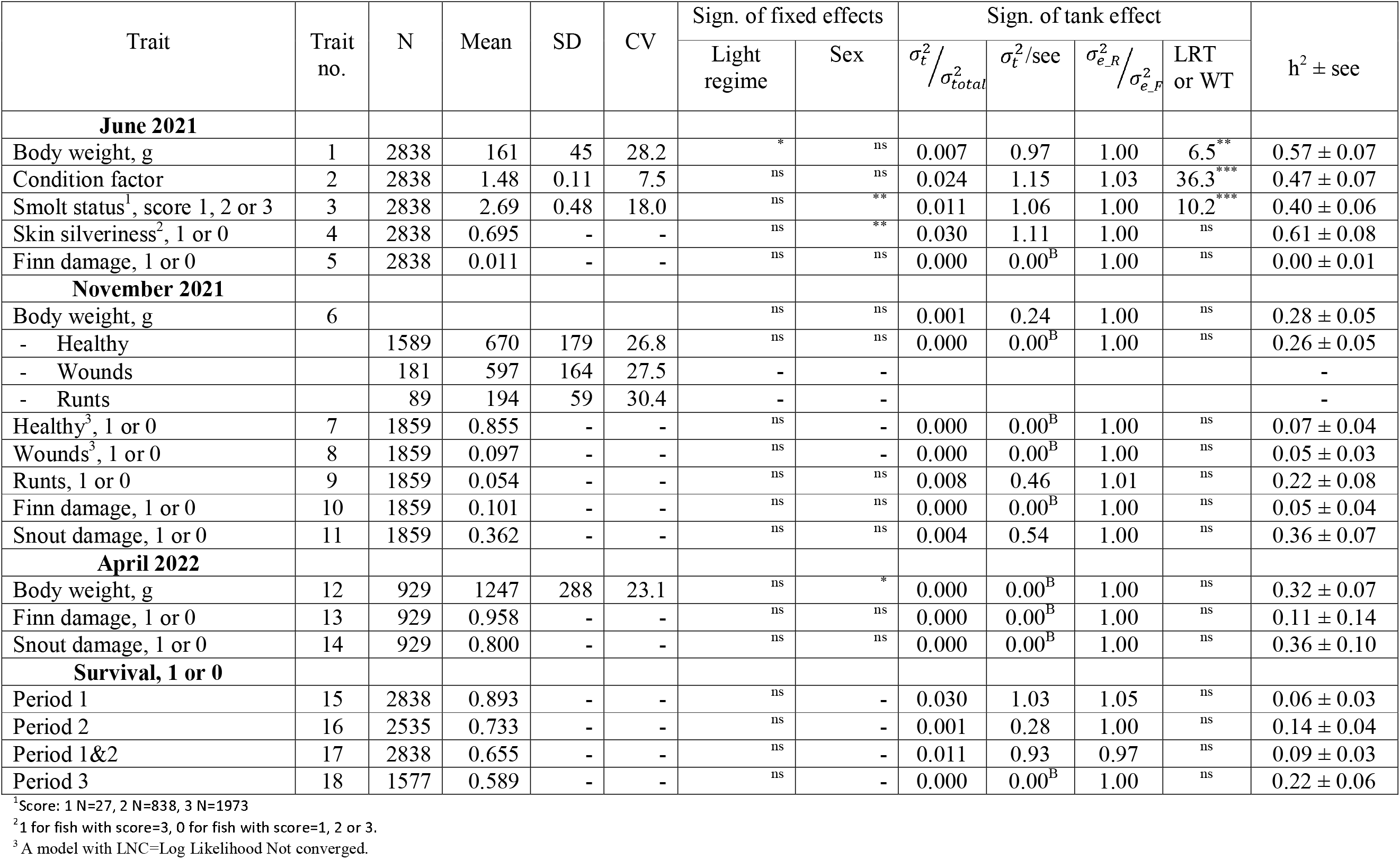
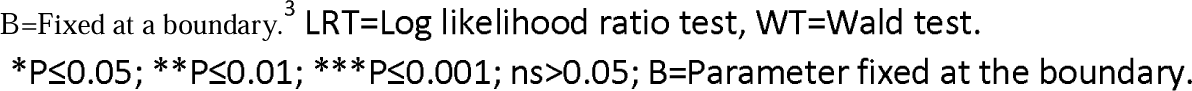
Descriptive statistics for each of the studied traits, the statistical significance of the two fixed effects and of the random tank effect, and the estimated heritability for each trait.

| Trait | Trait no. | N | Mean | SD | CV | Sign. of fixed effects |  | Sign. of tank effect |  |  |  | h<sup>2</sup> ± see |
| --- | --- | --- | --- | --- | --- | --- | --- | --- | --- | --- | --- | --- |
| | | | | | | Light regime | Sex | $\sigma_t^2 / \sigma_{total}^2$ | $\sigma_t^2$/see | $\sigma_{e\_R}^2 / \sigma_{e\_F}^2$ | LRT or WT | |
| June 2021 |  |  |  |  |  |  |  |  |  |  |  |  |
| Body weight, g | 1 | 2838 | 161 | 45 | 28.2 | * | ns | 0.007 | 0.97 | 1.00 | 6.5** | 0.57 ± 0.07 |
| Condition factor | 2 | 2838 | 1.48 | 0.11 | 7.5 | ns | ns | 0.024 | 1.15 | 1.03 | 36.3*** | 0.47 ± 0.07 |
| Smolt status<sup>1</sup>, score 1, 2 or 3 | 3 | 2838 | 2.69 | 0.48 | 18.0 | ns | ** | 0.011 | 1.06 | 1.00 | 10.2*** | 0.40 ± 0.06 |
| Skin silveriness<sup>2</sup>, 1 or 0 | 4 | 2838 | 0.695 | - | - | ns | ** | 0.030 | 1.11 | 1.00 | ns | 0.61 ± 0.08 |
| Finn damage, 1 or 0 | 5 | 2838 | 0.011 | - | - | ns | ns | 0.000 | 0.00<sup>B</sup> | 1.00 | ns | 0.00 ± 0.01 |
| November 2021 |  |  |  |  |  |  |  |  |  |  |  |  |
| Body weight, g | 6 |  |  |  |  | ns | ns | 0.001 | 0.24 | 1.00 | ns | 0.28 ± 0.05 |
| - Healthy |  | 1589 | 670 | 179 | 26.8 | ns | ns | 0.000 | 0.00<sup>B</sup> | 1.00 | ns | 0.26 ± 0.05 |
| - Wounds |  | 181 | 597 | 164 | 27.5 | - | - |  |  |  |  | - |
| - Runts |  | 89 | 194 | 59 | 30.4 | - | - |  |  |  |  | - |
| Healthy<sup>3</sup>, 1 or 0 | 7 | 1859 | 0.855 | - | - | ns | - | 0.000 | 0.00<sup>B</sup> | 1.00 | ns | 0.07 ± 0.04 |
| Wounds<sup>3</sup>, 1 or 0 | 8 | 1859 | 0.097 | - | - | ns | - | 0.000 | 0.00<sup>B</sup> | 1.00 | ns | 0.05 ± 0.03 |
| Runts, 1 or 0 | 9 | 1859 | 0.054 | - | - | ns | ns | 0.008 | 0.46 | 1.01 | ns | 0.22 ± 0.08 |
| Finn damage, 1 or 0 | 10 | 1859 | 0.101 | - | - | ns | ns | 0.000 | 0.00<sup>B</sup> | 1.00 | ns | 0.05 ± 0.04 |
| Snout damage, 1 or 0 | 11 | 1859 | 0.362 | - | - | ns | ns | 0.004 | 0.54 | 1.00 | ns | 0.36 ± 0.07 |
| April 2022 |  |  |  |  |  |  |  |  |  |  |  |  |
| Body weight, g | 12 | 929 | 1247 | 288 | 23.1 | ns | * | 0.000 | 0.00<sup>B</sup> | 1.00 | ns | 0.32 ± 0.07 |
| Finn damage, 1 or 0 | 13 | 929 | 0.958 | - | - | ns | ns | 0.000 | 0.00<sup>B</sup> | 1.00 | ns | 0.11 ± 0.14 |
| Snout damage, 1 or 0 | 14 | 929 | 0.800 | - | - | ns | ns | 0.000 | 0.00<sup>B</sup> | 1.00 | ns | 0.36 ± 0.10 |
| Survival, 1 or 0 |  |  |  |  |  |  |  |  |  |  |  |  |
| Period 1 | 15 | 2838 | 0.893 | - | - | ns | - | 0.030 | 1.03 | 1.05 | ns | 0.06 ± 0.03 |
| Period 2 | 16 | 2535 | 0.733 | - | - | ns | - | 0.001 | 0.28 | 1.00 | ns | 0.14 ± 0.04 |
| Period 1&2 | 17 | 2838 | 0.655 | - | - | ns | - | 0.011 | 0.93 | 0.97 | ns | 0.09 ± 0.03 |
| Period 3 | 18 | 1577 | 0.589 | - | - | ns | - | 0.000 | 0.00<sup>B</sup> | 1.00 | ns | 0.22 ± 0.06 |

<sup>1</sup>Score: 1 N=27, 2 N=838, 3 N=1973
<sup>2</sup>1 for fish with score=3, 0 for fish with score=1, 2 or 3.
<sup>3</sup>A model with LNC=Log Likelihood Not converged.
B=Fixed at a boundary.<sup>3</sup> LRT=Log likelihood ratio test, WT=Wald test.
\*P≤0.05; \*\*P≤0.01; \*\*\*P≤0.001; ns>0.05; B=Parameter fixed at the boundary.

### 2.3. eawater phase

On 23.06.2021, 2910 1+ smolts were transferred in a transport tank to the seawater facility of ARST. Fish from the three light treatments groups were kept in one 5.5 x 5.5 x 5.5 m sea net-cage and fed extruded salmon feed pellets (Supreme, Skretting AS, pellet size adjusted according to fish growth) until the end of the experiment in April 2022.

Eight weeks after seawater transfer, fish started to show signs of skin ulcers. Tissue from fish sampled on 25.08.2021 showed presence of *Moritella vicosa* that can cause skin ulcers and *Vibrio wodanis* (report from Norwegian Veterinary Institute, 9480 Harstad) often found in ulcers (Sommerset et al., 2022). The condition worsened and mortality rate increased over the autumn. The prevalence of ulcers persisted throughout the winter, despite removal of fish with wounds during the trait recordings of all fish in early November.

Mortality was monitored daily, and dead and moribund fish were removed from the sea cage and the moribund fish euthanized by a lethal dose of benzocaine (150 ppm). Dead and euthanised fish were identified with a PIT-tag reader, inspected visually for possible cause of death, after which their individual fork length and body weight were recorded.

Due to the high loss of fish and the associated welfare issues of the alive fish, it was decided to end the experiment in April 2022, approximately six months ahead of schedule.

The traits recorded during the November 2021 and April 2022 recordings are shown in Table 1. The thermal growth coefficient, as a measure of the overall growth rate was calculated as 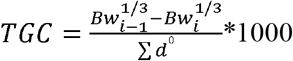 where *Bw* is the body weight at the start (*i*) and the end (*i-1*) of the actual period and the denominator 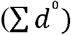 is the sum day degrees of the seawater temperature over the period (Jobling, 2003). TGC was calculated for the period between stocking (June 2021) and the November 2021 recordings, as well as between the November 2021 recording and the final April 2022 recording.

#### 2.3.1. November 2021 recordings

On 2-5 November 2022 batches of 20-30 fish were removed from the net pen with a dip-net and were anaesthetized in a seawater bath with benzocaine (50 ppm). The PIT-tag of each fish was recorded along with their body weight, fork length and health status (healthy, wounds, fin damage, snout damage and looser fish (with body weight < 300 g). Fish considered as healthy were put back into the cage.

#### 2.3.2. April 2022 recordings

On 19-22 April 2022 batches of 30 fish were removed from the net pen with a dip-net and were euthanized in a seawater bath with a lethal dose of benzocaine (150 ppm) until gill ventilation had ceased. The PIT-tag of each fish was recorded along with their body weight, fork length and health status (fin damage and snout damage).

#### 2.3.3. Survival in the seawater period

Three survival traits were defined based on whether a fish with recorded PIT-tag was recorded as dead (coded as 0) during three different periods in the net-cage in the sea, or alive (coded as 1) at the start or the end of the period.

- Period 1: Recorded as dead from day 0 (23.06.2021) to 58 (20.08.2021), otherwise coded as alive at day 0 (This is the period before the ulcers started).
- Period 2: Recorded as dead during day 59 to 135 (5.11.2021), otherwise coded as alive at day 58.
- Period 1&2: Coded as alive at day 135, otherwise coded as dead (but not necessarily recorded as dead) during day 0 to 135.
- Period 3: Coded as alive at day 300 (19.04.2022), otherwise coded as dead (but not necessarily recorded as dead) during day 136 to 300.

### 2.4. Statistical analyses

#### 2.4.1. Importance of the fixed effects and the random tank effect

The importance of the actual fixed (light regime and sex) and random (animal or sire and dam genetic) effects on the 16 studied traits (Table 1) was performed using bivariate mixed models in the ASReml software (Gilmour et al., 2021). This software was also used also for the other statistical models.

First, a bivariate linear mixed animal models was used for body weight in June 2021 (*y*_1_) and for each of the other continuous traits (*y*_2_= trait 2 (condition factor), 3 (smolt status), 6 (body weight recorded in November 2021) or 12 (body weight recorded in April 2022), one at a time) in Table 1:

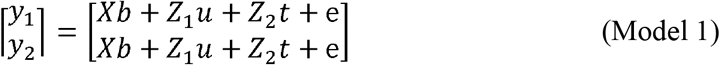

Where ***y***_**1**_ is a column vector for body weight in June and ***y***_**2**_ is a column vector for trait 2, 3, 6 or 10; ***X*** is an incidence matrices that assign each trait record to the appropriate level of the fixed light regime and sex; ***Z***_**1**_ and ***Z***_**2**_ are incidence matrices that assign each trait record to the appropriate level of the random effects of animal and tank, respectively; ***b*** is a vector of the fixed effects of light regime (with three levels) and sex (with two levels) determined by genetic markers located within the sdY gene (Houston et al., 2014); ***u*** is a vector of random additive genetic values for animal with 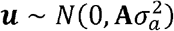 where 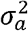 is the additive genetic variance and ***A*** is the additive genetic numerator relationship matrix; ***t*** is the vector of random effect of tank with two levels within each of the three light regimes with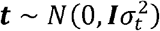, where 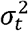 is the tank variance, and ***e*** is a vector of random residual effects with 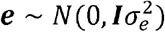 where 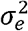 is the residual variance.

For the binary traits in Table 1 a linear mixed sire and dam model was used for body weight in June 2021 as one of the trait (*y*_1_) and a mixed sire and dam threshold model for each of the binary traits (*y*_2_); i.e. trait 4, 5, 7, 8, 9, 10, 11, 13, 14, 15. 16, 17 and 18, one at a time:

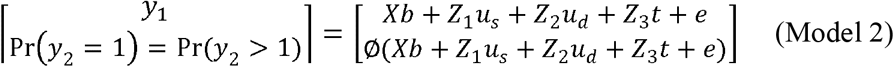

where for the binary trait Φ(·) is the cumulative standard normal distribution function assuming a normal underlying liability variable, *l*, determining the categorical outcome of the binary trait such that *l* ≤0 corresponds with *y*_2_=0, and *l* >0 with *y*_2_=1); ***u***_s_ and ***u***_d_ are random_s_ additive genetic values for sires and dams, respectively with 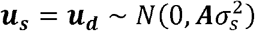 and 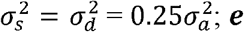 is a vector of random residual effect with 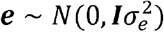 and 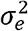 is the residual variance including half of the additive genetic or the Mendelian sampling variance; and the other parameters and matrices are as described for Model 1. For each of the bivariate traits 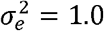. Body weight in June 2021 was included as a trait to account for possible bias in the estimates due to mortality that may be genetically correlated to each of the other traits.

The importance of the effect of light regime was tested using a Wald F-test statistics with two (numerator) and three (denominator) degree of freedom.

For each of the studied traits the importance of the random tank effect was expressed as (a) the ratio of its estimated variance component relative to the total variance 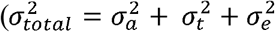 for the continuous traits and 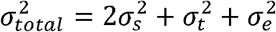 for the binary traits), (b) the ratio of its estimated variance component and its standard error (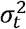; an output in the ^*^asr file in ASREML), (c) the ratio of the estimated variance component of the error of the reduced model (after omitting the tank effect) and the full model (for each of the continuous traits) and as the ratio of the estimated variance component of the sire effect of the reduced model (after omitting the tank effect from the model) and the full model (for each of the binary traits as the error variance for each of the binary traits was set equal to unity both in the full and the reduced model).

In addition, the importance of the random tank effect for each of the continuous traits was tested using a log likelihood ratio test (LRT, *χ*^*2*^ variate with 1 degree of freedom, Wald, 1943; Lynch and Walsh, 1997) at the lower boundary of its parameter space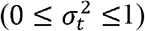; thus using a 5% critical value of 2.71 in contrast to 3.84 for a *χ*^*2*^variate with 1 degree of freedom (see chapter 2.4.1 in Gilmore et al. 2021). REML estimates of variance components cannot generally be compared with LRT tests because they are not true maximum likelihood (ML) estimates. However, in practice parameter estimates for continuous traits from ML and REML produce very similar estimates from large samples as is the case for the present data. For the binary traits the importance of the random tank effect was tested using a Wald Chi-Square Test (WT, *χ*^*2*^ variate with 1 degree of freedom; Wald, 1943?) as LRT is not an appropriate test statistic for binary traits.

For the binary traits the least squares estimate on the liability scale was transformed back to observed binary scale.

Included in the above models was also an additional random effect common to fullsibs other than additive genetics, i.e., non-additive genetic effects. However, for all traits found this effect was to be not significantly different from zero and was therefore omitted from the above models, as well as for the below models.

#### 2.4.2. The same trait at the different light regimes as different traits

Separately for each of the three recordings (June 2021, November 2021 and April 2022), estimates of genetic correlations between the same trait for fish that received different light regimes were obtained from a three-trait linear mixed animal model for the each of continuous traits (Model 3) and from a mixed sire and dam threshold model for each of the binary traits (Model 4):

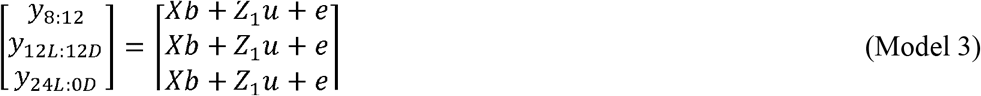

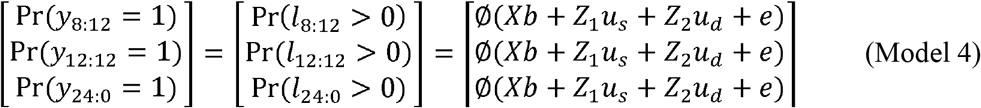

where ***y*** for each of the three light regimes is a column vector of the actual trait at the June 2021, Nov 2021 or the April 2022 recording; and the other parameters and matrices are as described for Model 1. For both Model 3 and 4 the residual correlations between the same trait at the three different light regimes were set equal to zero as the trait was recorded on different animals. For the traits recorded in November 2021 and April 2022, body weight in June 2021 was included as a fourth trait to account for possible bias in the estimates due to mortality that may be genetically correlated to the other traits.

For some of the traits the log likelihood of the model did not converge, or the log likelihood converged while the parameters did not converge; in most cases due to one, two or all three of the estimates of the genetic correlations at the upper boundary (unity) of the parameter space. For those cases one or two of the three genetic correlations were fixed close to the upper boundary (0.99) and at which both the log likelihood and the parameters converged, in most of these cases.

#### 2.4.3. Estimates of genetic parameters

For the traits in Model 3 for which the log-likelihood and the parameters converged and with a heritability significantly different from zero, the genetic correlations between the same trait for fish that received different light regimes were found to be high or very high (Table 2).

**Table 2.** Estimates of genetic correlations between the same trait at the different light regimes^1^.

| Trait | Time | LR1 - LR2 | LR1 - LR3 | LR2 - LR3 | Comment<sup>2</sup> |
| --- | --- | --- | --- | --- | --- |
| Body weight | June 2021 | 0.96 ± 0.04 | 0.99<sup>B</sup> | 0.97<sup>F</sup> | LLC, PC |
|  | Nov. 2021 | 0.92<sup>F</sup> | 0.96 ± 0.08 | 0.96 ± 0.08 | LLC, PC |
|  | April 2022 | 0.96<sup>F</sup> | 0.87 ± 0.15 | 0.74 ± 0.16 | LLC, PC |
| Cond. factor | June 2021 | 0.99<sup>F</sup> | 0.89 ± 0.06 | 0.98<sup>7</sup> ± 0.04 | LLC, PC |
| Smolt status | June 2021 | 0.99 ± 0.05 | 0.98 ± 0.05 | 0.99<sup>F</sup> | LLC, PC |
| Skin silveriness | June 2021 | 0.99<sup>B</sup> | 0.99<sup>B</sup> | 0.99<sup>B</sup> | LLC, PC |
| Fin damage | Jun. 2021 | - | - | - | CSNPD |
|  | Nov. 2021 | - | - | - | CSNPD |
|  | April 2022 | 0.99<sup>F</sup> | 0.99<sup>B</sup> | 0.99<sup>B</sup> | LLC, PC |
| Snout damage | Nov. 2021 | 0.99<sup>F</sup> | 0.99<sup>B</sup> | 0.89 ± 0.20 | LLC, PC |
|  | April 2022 | 0.99<sup>B</sup> | 0.99<sup>B</sup> | 0.99<sup>B</sup> | LLC, PC |
| Healthy | Nov. 2021 | - | - | - | CSNPD |
| Wounds | Nov. 2021 | - | - | - | CSNPD |
| Runts | Nov. 2021 | - | - | - | CSNPD |
| Survival | Period 1 | - | - | - | CSNPD |
|  | Period 2 | 0.99<sup>F</sup> | 0.86 ± 0.38 | 0.73 ± 0.35 | LLNC |
|  | Period1&2 | 0.81 ± 0.33 | 0.47 ± 0.43 | 0.65 ± 0.37 | LLC, PC |
|  | Period 3 | 0.84<sup>P</sup> ± 0.35 | 0.83<sup>P</sup> ± 0.31 | 0.41<sup>P</sup> ± 0.26 | LLNC |

<sup>1</sup>LR1=8L:16D; LR2=12L:12D; LR3=24L:0D;
<sup>2</sup>LLC=Log Likelihood Converged;
PC=Parameters converged;
CSNPD=Correlation Structure Not Positive Definite;
LLNC=Log Likelihood Not converged.
Superscript : F=Fixed by user to the given value; B=Fixed at a boundary; P=Positive definite.

Consequently, for these traits the same trait for the three different light regimes at each of the three recordings could be looked upon as the same trait, thus reducing the traits 1 to 18 in Table 1 to eight traits: body weight, condition factor, smolt status (or skin silveriness that showed a unity genetic correlation to smolt status) in June 2021, body weight, runts and snout damage in November 2021, and body weight and snout damage in April 2022.

Estimates of (co)variances among these traits were obtained from the following multitrait mixed animal model:

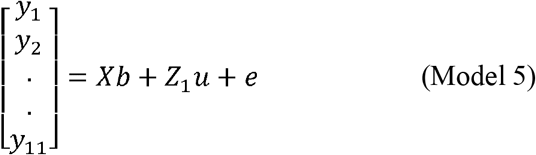

where *y*_1_, *y*_2_*…, y*_11_ are vectors of the animal trait records for each of the twelve traits; ***X*** is an incidence matrices that assign each trait record to the appropriate level of the combined fixed effect of light regime and tank and of the fixed effect of sex; ***b*** is a vector of the combined fixed effects of light regime and tank, and the fixed effect of sex; and the other matrices and parameters are as described for Model 1. When all eight traits were included in the model neither the log likelihood nor the parameters did not converge, but converged after omitting the runts trait in November 2021; thus remaining with the seven traits in Table 3.

**Table 3.** Estimates of genetic (above diagonal) and residual^1^ (below diagonal) correlations between the seven traits from Model 2.

| Time recorded | Trait | June 2021 |  |  | November 2021 |  | April 2022 |  |
| --- | --- | --- | --- | --- | --- | --- | --- | --- |
|  |  | Body weight | Cond. factor | Smolt status | Body weight | Snout dam. | Body weight | Snout dam. |
| June 2021 | Body weight | | $-0.12 \pm 0.15$ | $0.68 \pm 0.15$ | $0.63 \pm 0.17$ | $-0.10 \pm 0.12$ | $0.65 \pm 0.12$ | $0.14 \pm 0.12$ |
| | Cond. factor | 0.29 | | $-0.35 \pm 0.17$ | $-0.28 \pm 0.16$ | $0.00 \pm 0.12$ | $-0.37 \pm 0.12$ | $0.43 \pm 0.12$ |
| | Smolt status | 0.21 | 0.07 | | $0.48 \pm 0.12$ | $-0.08 \pm 0.18$ | $0.61 \pm 0.18$ | $0.02 \pm 0.14$ |
| Nov. 2021 | Body weight | 0.38 | 0.05 | 0.12 | | $0.00 \pm 0.19$ | $0.86 \pm 0.19$ | $0.12 \pm 0.15$ |
| | Snout damage | 0.18 | 0.04 | 0.09 | 0.21 | | $-0.22 \pm 0.20$ | $0.81 \pm 0.16$ |
| April 2022 | Body weight | 0.39 | 0.10 | 0.09 | 0.81 | 0.13 | | $-0.12 \pm 0.12$ |
| | Snout damage | 0.14 | -0.08 | 0.10 | 0.34 | 0.08 | $0.22 \pm 0.12$ | |

<sup>1</sup> Standard errors of all residual correlations were $< 0.01$ .

**Table 4.**
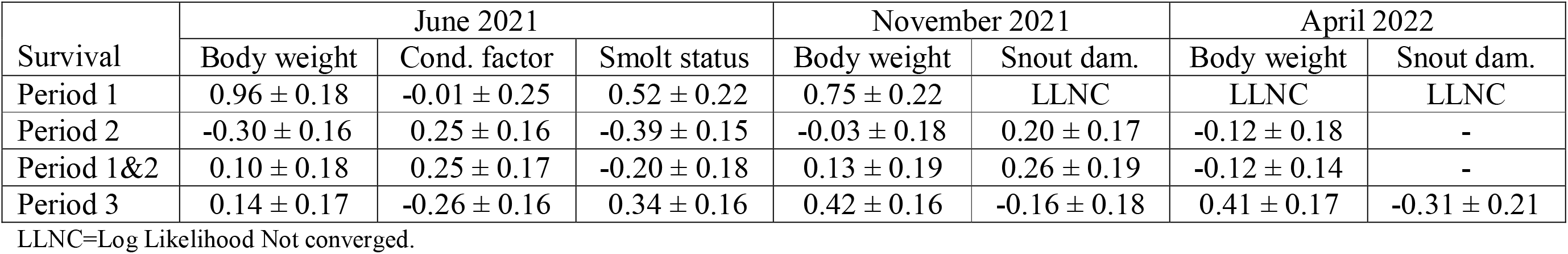
Estimates of genetic correlations of the survival traits with the seven traits in Table 3.

For the four survival traits in Table 1, covariances of each of these traits with the seven traits that converged in Model 5, were obtained from multi-trait sire and dam models similar to Model 2 with the above-mentioned binary traits included in the model, one at a time, in addition to the seven traits in Model 5.

For the traits in Model 2 the sire component of variance was set equal to the dam component using the model function and(dam,1) in the ASREML software. For the binary traits on the underlying liability scale the residual variance 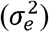 was set equal to unity and hence only variance components for sire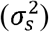 and full-sib 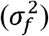 need to be estimated.

Hence, for the animal model (Mode 2) the heritability for each trait was calculated as 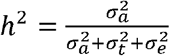, and for the sire and dam models as 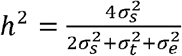.

The relative magnitude of the tank effect was expressed as 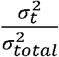 where the total variance in the denominator is 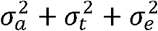 for the continous traits and 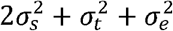 for the binary traits.

## 3. Results

### 3.1 Overall means

Table 1 shows the number of animal trait records at each of the three recordings in June 2021, November 2021 and April 2022. At the June recording, 69.5 % of the fish obtained the highest score for smolt status. The proportion of fish with fin damage was very low (1.1%), low (10.1%) and high (95.8%) at the June 2021, Nov. 2021 and April 2022 recordings, respectively. At the November 2021 recording 9.7% of the fish had wounds, 5.4% were classified as runts and the remaining 84.8% as healthy. The proportion of fish with snout damage was moderate at the November 2021 recording (36.2 %) and high (80.0 %) at the April 2022 recording.

Based on the difference between the number of fish at the start and the end of each of the three growth periods, the overall survival was 89.3% in Period 1 (day 0 to 58), declining to 73.3% in Period 2 (day 59 to135) and to 58.9% in Period 3 (day 136 to 300). The overall survival in the entire seawater period was 34.8% (929/(2938-181-89).

However, the percentages of dead recorded fish and with readable PIT-tags were lower as not all dead fish could be found in the cage. In Period 1&2, 778 fish were recorded as dead and with readable PIT-tags; i.e. 79.5 % (778/(2838-1859) of the difference between the number of alive fish at the recordings in June 2021 and November 2021. In Period 3, 363 fish were recorded as dead and with readable PIT-tag; i.e. 56.0 % (363/(1577-929) of the difference between the number of healthy fish at the November 2021 and the number of alive fish at the April 2022 recording. For the entire seawater period 70.0% (778+363)/(2838-181-89-929) of the dead fish were recorded as dead with readable PIT-tag. Consequently, as date of death could not be obtained for 29.9% of the dead fish, the number of days to death was not considered as a relevant trait in this study.

Based on the dead recorded fish with PIT-tags survival curves in the seawater period for each of the three light regimes are shown in Figure 1. Fish in group 24L:0D showed increased mortality from day two after sea transfer with an accumulated mortality of 8.9% during Period 1 and 1.0% for the 12L:12D group and 0.7% for the 8L:16D group. From day 59 onwards until harvest at day 300 in April the mortality was very similar for the fish on the three light regime groups. Worth a notice is that these survival curves are grossly overestimated as all dead fish could not be found in the cage.

**Figure 1.**
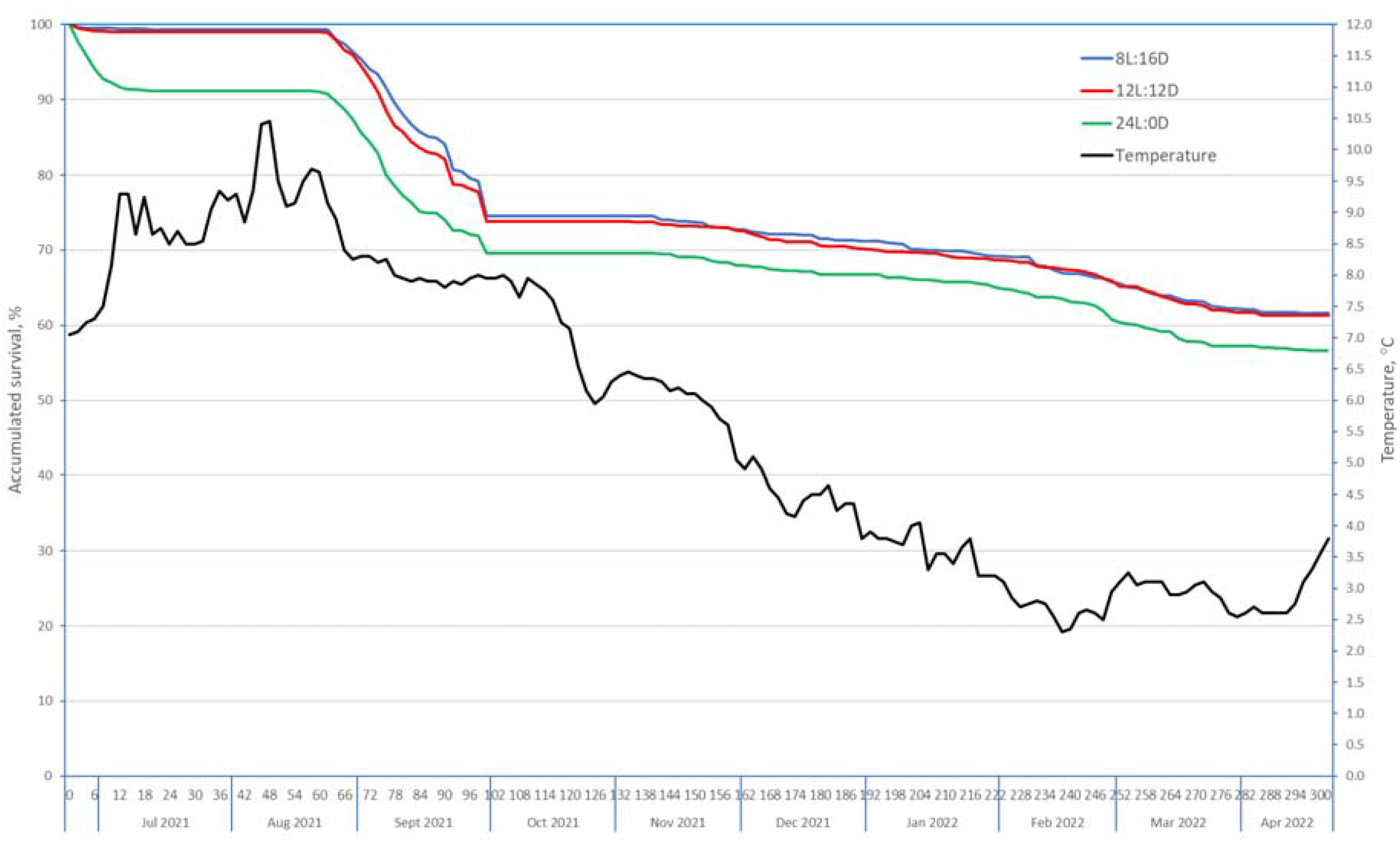
Survival (%) curves in the net-cage in the sea for fish that received three different light regimes during the smoltification period (based on the fish that were recorded as dead and with readable PIT-tag).

For fish with recorded body weights TGC was 2.80 (SD=0.85) in Period 1&2 and 3.02 (SD=0.89) in Period 3. For fish recorded as healthy at the Nov. 2021 recording TGC in Period 1&2 was 2.97 (SD=0.64), 2.61 (SD=0.60) for fish classified as runts, and 0.62 (SD=1.01) for fish classified with wounds.

### 3.2 Effect of light regime

Body weight in June 2021 was the only trait significantly affected by light regime (Table 1) with fish on 24L:0D being 15.0 ± 4.0 (10.0%) and 18.0 ± 4.0 (12.2%) gram heavier than fish on light regime 12L:12D and 8L:16D, respectively (Figure 2).

**Figure 2.**
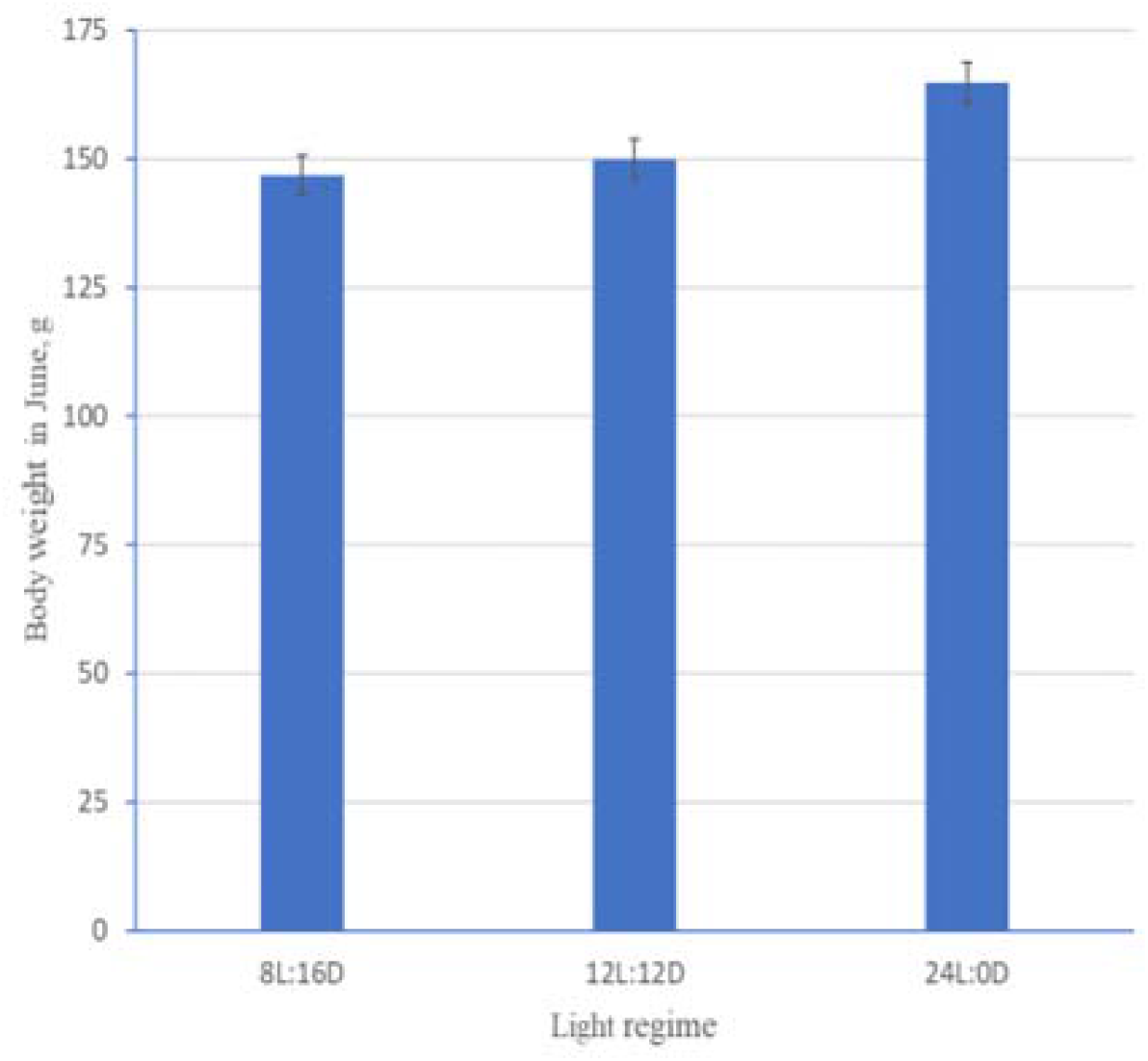
Least square means estimates (±see) for body weight in June 2021 for the three light regimes.

For survival in Period 1 the estimates threshold on the liability scale was 0.985 ± 0.142 for fish on light regime 24L:0D. This threshold was lower than for fish on light regime 12L:12D (-0.397 ± 0.191 and 8L:16D (-0.350 ± 0.191), corresponding to a difference in survival of -8.3 and -7.5 %-units on the observed scale, as compared to -8.0 and -8.3 %-units based on the data in Figure 1 which do not include the non-recorded dead fish. However, these above threshold differences were not significantly different from zero judged by the Wald F-test statistics for this effect in Model 2 (F=2.55, critical F-value 9.55 for P=0.05).

Figure 3 shows that prior to seawater transfer the survivors of all three light regimes in Period 1, Period 1&2 and Period 3 had more favorable values for the smolt indicator traits; i.e. lower condition factor, higher smolt status score and higher skin silveriness proportion than the fish that died in these periods. In addition, the survivors of all three regimes had a higher body weight in Period 1 and Period 3, but not in Period 1&2. Significant differences between light regimes for the difference between the survivors and the dead were observed for all traits in Period 1, and for condition factor in Period 1&2. In Period 3 no significant difference between the survivors and the dead between light regimes was found for any of the four traits recorded prior to seawater transfer. Although most of the estimated differences between the survivors and the dead were small relative to their actual trait means and with large standard error, they indicate that fish that had received a shorter light day for six weeks until six weeks prior to seawater transfer (8L:16D or 12L:12D) were at a more optimal smolt status at the time of seawater transfer than fish that had been kept on a continuous light regime (24L:0D). Between light regimes the following significantly differences between survivors and the dead were found:

**Figure 3.**
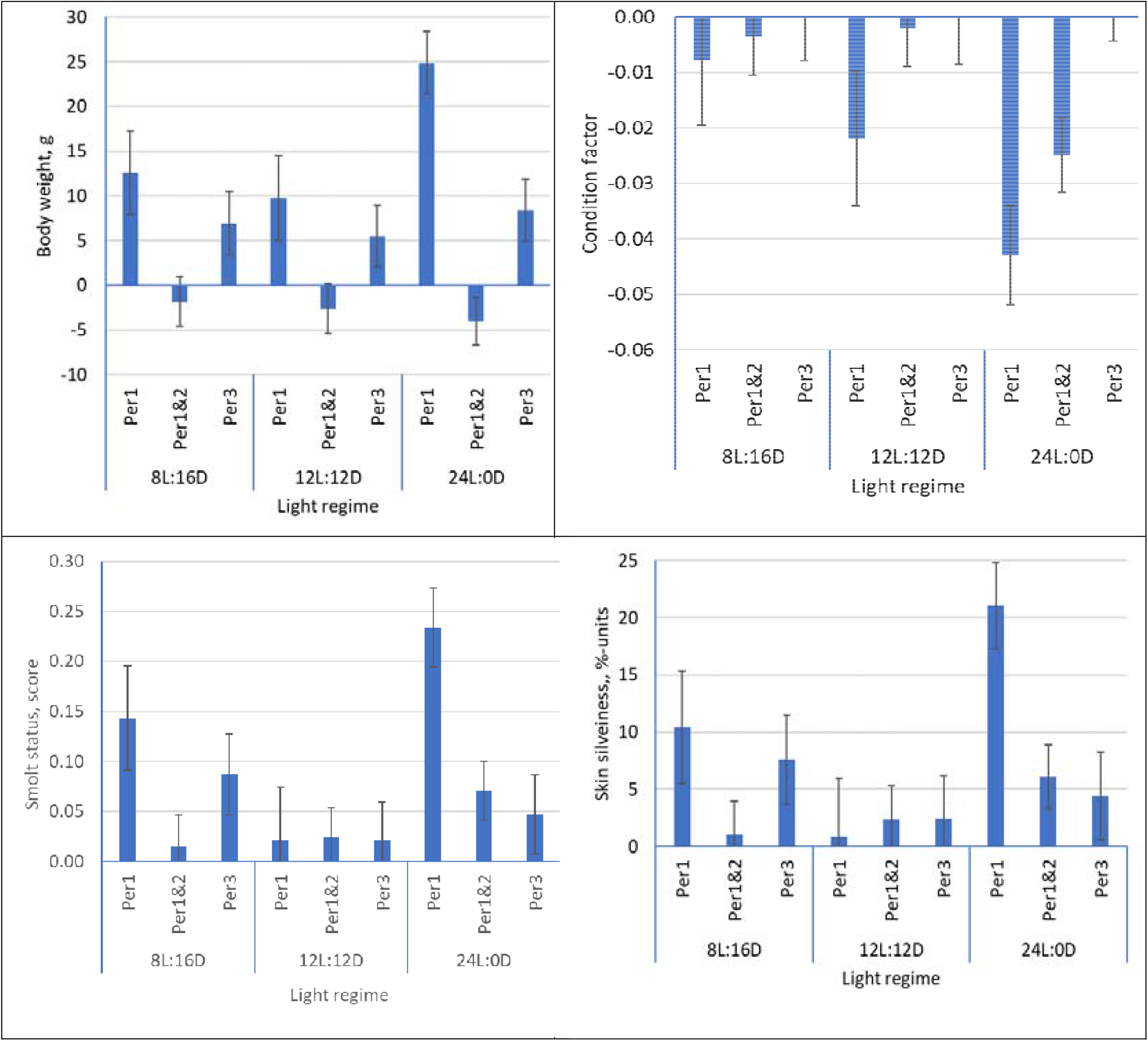
Estimates of the difference (±see) between the survivors and the dead fish in Period 1, Period 1&2 and Period 3 for traits recorded in June 2021 for each of the three light regimes.

For condition factor in Period 1, the estimated difference between survivors and the dead was significantly larger (negative) for fish on the 24L:0D than for fish on the 8L:16D (P=0.017) regime, while in Period 1&2 the estimated difference was significantly larger for fish on the 24L:0D regime than for fish on the 8L:16D (P=0.026) and on the 12L:12D (P<0.017) regimes.

For smolt status score in Period 1 the estimated difference between the survivors and the dead was significantly lower for fish on the 24L:0D than for fish on the 12L:12D (P=0.001) regime. For skin silveriness in Period 1 the estimated difference between the survivors and the dead was significantly higher for fish on the 24L:0D than for fish on the 8L:16D (P=0.002) regime.

For body weight in Period 1 the estimated difference between the survivors and the dead was significantly larger for fish on the 24L:0D than on the 8L:16D (P=0.035) and the 12L:12D (P<0.011) regimes.

### 3.3 Sex

Table 1 shows that sex had a significant effect on only three of the studied traits, i.e., smolt status score and skin silveriness in June 2021 and body weight in April 2022. In June 2021 the males had a significantly higher score for smolt status (0.044 ± 0.016, 1.6%; P<0.01), a significantly higher liability threshold for skin silveriness (0.151 ± 0.050, 4.8 %-units on the observed scale; P<0.01), and in April 2022 a significantly higher body weight (34.9 ± 15.8 g or 2.9%; P<0.05).

### 3.4 Tank within light regime

Body weight, condition factor and smolt status score in June 2021 were the only traits for which the tank effect was significantly different from zero (P<0.05) (Table 1). Of the total variance of the trait it accounted for 0.7% for body weight, 2.4 % for condition factor and 1.1% for smolt status score. For most of the other traits the tank effect accounted for an even smaller proportion of the total variance. However, for the two binary traits skin silveriness in June and survival in Period 1 the tank effect accounted for 3.0% of the total variance.

For survival in Period 1 the magnitude of the tank effect is illustrated in Figure 4 where the estimated tank solution for each of the six tanks are shown both on the liability scale and the back transformed observed scale. The power of detecting the observed survival difference in Period 1 between the 24L:0D regime (83.4%) and the mean survival of the 8L:16D and 12L:12D regimes (0.5(90.9+91.6)=91.3%) was found to be 0.41 only (two-sided test with a 0.05 Type-1 error); calculated on the observed binary scale based on the estimated variance component of the effect of tank within light regime as a proportion of the sum of the variance component of the tank effect and the variance component within tank (0.001569/(0.001569+0.092576)=0.0167).

**Figure 4.**
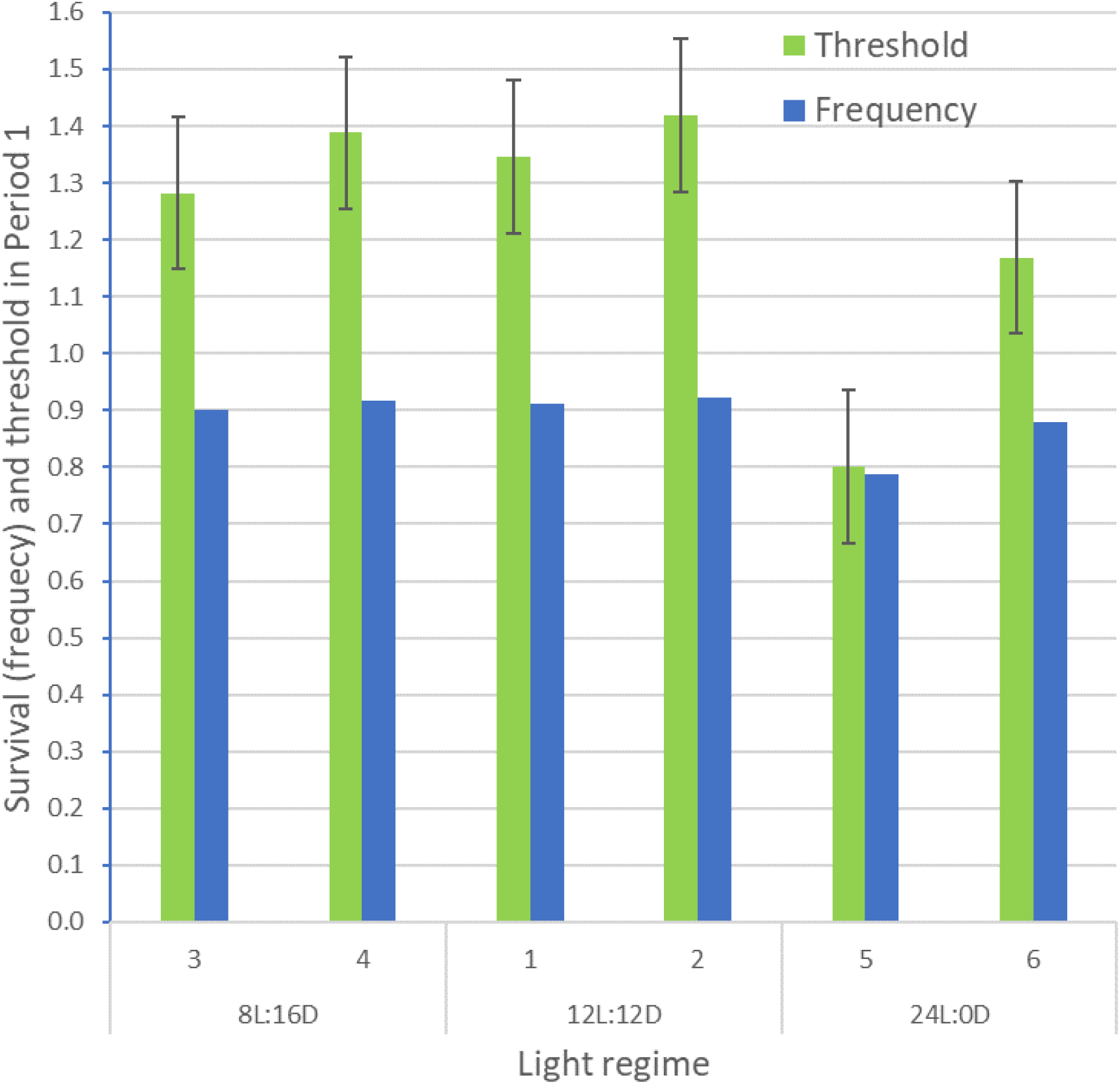
The estimated liability thresholds for survival in Period 1 for each of the two tanks per light regime and their corresponding back transformed survival frequency values.

If the number replicated tanks per light regime was increased to 4, 6 or 8 and with the same total number of fish per treatment, the power would increase to 0.64, 0.77 and 0.85, respectively. These results clearly illustrate that in this experiment the number of replicated tanks per light regime should have been at least six to detect a 7.9%-units difference in survival between two light regimes with a sufficiently high power.

If the magnitude of the replicated tank effect could have been reduced to 0.01, 0.005 or 0.0025 of the total variance, the power of detecting a 7.9%-units difference in survival between two light regimes would be 0.58, 0.81 and 0.94, respectively with two replicated tanks per light regime; and 0.81, 0.94 and 0.98, respectively with four replicated tanks per light regime.

### 3.5 The same trait at the different light regimes as different traits

For 12 of the 18 studied traits the genetic correlations between the same trait at the three different light regimes were generally high to very high (Table 2). However, for six (all binary traits) of the 18 traits the correlation structure was not positive definite (Table 2).

### 3.6 Heritability

The heritability estimates for the external indicator traits for smolt development (condition factor, smolt status score and skin silveriness) were moderate, as were the heritability estimates for snout damage both in November 2021 and April 2022, runts in November 2021 and Survival in Period 3 (Table 1). The heritability estimates for fin damage in both June 2021, November 2021 and April 2022, for healthy fish and wounds in November, and for survival in Period 1 and Period1&2 were all low (Table 1).

### 3.7 Genetic and residual correlations among the traits

Estimates of genetic and residual correlations among the seven traits included in Model 2 are given in Table 3. The genetic correlations of smolt status in June 2021 with body weight in June 2021, November 2021 and April 2022 were all moderately positive and thus favorable. The genetic correlation between snout damage in Nov. 2021 and April 2022 was relatively high (0.81), while the genetic correlation of snout damage with the other traits were in general low and not significantly different from zero with one exception; that between condition factor in June 2021 and snout damage in April 2022 (0.43). The genetic correlation between skin silveriness (%) and smolt status (score) was close to unity (fixed by the program at 0.9985 and thus close to the upper boundary).

The genetic and residual correlations between body weights in June, November and April were all positive, and those of June with November 2021 (r_g_=0.63; r_e_=0.38) and April 2022 (r_g_=0.65; r_e_=0.39) lower than those between November and April (r_g_=0.86; r_e_=0.81). The magnitude of the correlations between these traits and of their heritability estimates (Table 1) are similar to earlier published estimates in Atlantic salmon (Gjerde et al., 1994).

The remaining genetic and residual correlations in Table 2 were all close to zero.

### 3.8 Genetic correlations of the survival traits with the other traits

Table 3 shows that the genetic correlation of body weight in June 2021 with survival in Period 1 was high (0.96), but close to and not significantly different from zero with survival in Period 1&2 and Period 3. The genetic correlations of survival in Period 1 with smolt status in June 2021 and body weight in November 2021 were positive but with large standard errors and thus not significantly different from zero (P>0.05). The genetic correlations of the survival traits with the other traits recorded in June 2021, November 2021 and April 2022 were all of low to medium in magnitude and with large standard errors and thus not significantly different from zero (P>0.05).

Estimates of the genetic correlation of the runts trait with the other traits could not be obtained as none of the (co)variance components converged.

## 4. Discussion

The objective of this study was to assess the impact of three different light regimes during 12 weeks prior to seawater transfer and fish genetics, as well as the interaction between light regimes and genetics, on the development of external smolt characters, survival in seawater, growth, and general fish welfare characteristics.

### Light treatment result in more synchronous smolt development

The three investigated light regimes impacted several smolt traits. In line with similar experiments (Striberny et al., 2021; Ytrestøyl et al., 2022) we found that during the light regimes, fish on the two short day smoltification regimes (8L:16D, 12L:12D) had slower growth compared to fish on the continuous light regime (24L:0D), likely a negative effect of shorter daylength on appetite (Bjørnsson et al., 1989). After 5 and 10 months in the sea, however, there was no longer a significant effect of light regime on fish size (P>0.05). This “catching up in sea” effect also agrees with previous studies (Striberny et al., 2021) showing that fish having slower growth during short day smoltification regimes but grow better in seawater compared to fish kept on continuous light.

For wild salmon, circannual changes in daylength are important cues that regulate the timing of smoltification and subsequent migration to sea (Hoar, 1988). Experimental daylength manipulations have shown that both simulated natural increase in daylength in spring and an abrupt increase in daylength to mimic the transition from a winter to a spring photoperiod impact hormonal regulation and development of smolt physiology and seawater adaptations (McCormick et al., 2007). It is therefore possible that fish raised on constant light in our study were less ‘synchronized’ in their smolt development and were more variable in their seawater adapted physiological states. This corresponds with several previous studies that have reported a lack of synchronization in smolt development of Atlantic salmon kept on long photoperiod (Björnsson et al. 1995, Sigholt et al. 1995, Duncan and Bromage 1998). Several results in this study support this notion. Firstly, over the first 58 days in the net-cage in the sea (Period 1) mortality was to a large degree confined to fish raised on the 24L:0D regime (Figure 1). Secondly, difference in traits measured prior to seawater transfer between fish that survived and those that died in Period 1 were largest in the 24L:0D group (Figure 3) which strongly indicates a positive effect of a short day smoltification regimes, not only on survival after seawater transfer, but also on the general synchronization (i.e., group level uniformity) of physiological processes during smoltification. Finally, the non-significant effect of light regime on the welfare traits (fin and snout damage and runts) recorded in November 2021 and April 2022 strongly indicates that a reduced daylight regimes can be recommended without a negative effect on these traits.

A positive effect of increased body size at seawater transfer on survival in both Period 1 and 3 was observed for all three light regimes and one of the reasons why smolt producers tend to produce a large smolts. However, the results from the study of Ytrestøyl at al. (2022) referred to in the introduction show that the choice of optimal smolt size and smoltifying strategy is not straight forward procedure.

The non-significant effect of light regime on survival in Period 1 (Table 1) might be considered strange given the much lower observed survival of the fish in the 24L:0D group than in the 8L:16D and 12L:12D groups. The main reason for this is the substantial although non-significant tank effect on this trait resulting in a low power of the experiment; i.e. low probability of rejecting the null hypothesis (no difference between the light regimes) when the alternative hypothesis of a significant difference is true. Therefore, in a similar future experiment, given a similar sized tank effect as in this study, the number of replicated tanks per light regime should be increased to at least six and thus to a total of 18 instead of six tanks; a relative marginal increase in the experimental cost as the fish are to be kept in these tanks during the first six weeks of the 12 weeks light regime, after which the fish from all three light regimes in the following six weeks could have been PIT-tagged and pooled and reared in a common tank on the 24L:0D regime; a strategy that may have reduced the magnitude of the tank effect and increases the power and thus the probability to detect also a smaller true difference for a trait between the light regimes.

### Can we breed a more robust smolt?

We found that condition factor, smolt status score and skin silveriness (a binary trait derived from the smolt status score trait), all considered as external indicator traits for smolt development, have a significant genetic component and thus can be improved through selection. A significant genetic variation in condition factor was also reported by Khaw et al. (2021), in both 0+ and 1+ smolts. The very high genetic correlation between each trait recorded on sibs exposed to the different light regimes shows that each trait at the different light regimes can be looked upon as the same trait. Very high genetic correlations were also found between the same trait of the three light regimes recorded both in November 2021 and in April 2022. Consequently, group of sibs from the same families smoltified under different light regimes are expected to rank very similar for these traits. It is therefore irrelevant under which light regime the test fish and the breeding candidates in a nucleus breeding population, or the commercially fish produced from selected breeders in a breeding program, are being smoltified. Therefore, selection for improved smolt status through selection for e.g., a lower condition factor and/or a higher smolt status score should result in a favourable selection response in the traits irrespective under which light regime the fish are being smoltified.

The above-mentioned positive effect of the magnitude of the traits recorded prior to seawater transfer on survival in Period 1 was also seen at the genetic level as a very high genetic correlation (0.96) between body weight in June 2021 and survival in Period 1, both within the continuous (24L:0D) and the short day (8L:16D and 12L:12D) light regimes. This strongly indicates that in a group of fish expected to be either well (8L:16D and 12L:12D) and poorly smoltified (24L:0D) those with a high genetic potential for growth have a higher probability to survive during the first weeks in seawater as compared to fish with a lower genetic growth potential.

The much higher estimated genetic correlation of survival in Period 1 with body weight (0.96) than with smolt status (0.52) and condition factor (-0.01) in June 2021 strongly indicate that growth rate until the time of seawater transfer is of greater importance for the probability to survive in the first weeks in the sea than the subjectively scored smolt status, and that condition factor in June is a very poor predictor for survival in the sea.

It is therefore possible to selectively breed for salmon with an overall more desired smolt characteristics (skin silveriness and condition factor). However, our growth and survival data in the sea-phase questions these traits as good predictors for smolt performance in the sea. Firstly, the differences in the mean values of the traits recorded prior to sea transfer between the survivors and the dead in Period 1 were relatively small thus indicating a substantial variation among the individuals in other unknown trait values of importance for the ability of the salmon to thrive in seawater. Secondly, genetic correlations of survival in seawater with condition factor and smolt status prior to sea-water transfer were in general low, meaning that direct selection for these smolt indicator traits is expected to result in a low correlated genetic response in survival in the seawater phase. However, the magnitude of these genetic correlations has high uncertainty as seen by their large standard errors. To what extent the above genetic parameters are also valid for fish being of an on average larger or smaller body size at the time of seawater transfer than the fish in this study needs a further study.

Therefore, indirect selection for higher survival in seawater through selection for a salmon that at seawater transfer have higher smolt status score or a lower condition factor is a questionable strategy as this would require an indicator trait with a much higher genetic correlation to the breeding objective trait survival than found in this study (Falconer and McKay, 1989). A most likely better strategy would be to perform direct selection for increased survival and growth during the first months in the sea but would require that the individual body weights of all fish were recorded just prior to or after being smoltified, and of all the survivors after e.g. three to four months after sea transfer. Use of genomic selection (Meuwissen, et al., 2016; Boudry et al. 2021) would increase the selection for particular the survival trait through both higher accuracy and intensity of selection.

The moderate heritability for snout damage (0.36 in both November and April) and the relatively high genetic correlation between this trait at the two recordings (0.81) strongly indicate that the trait has a substantial genetic component. Consequently, its frequency that was substantial both in November (36.2%) and April (80.0%) can thus be reduced through direct selection against the trait which is not expected to take place through a correlated response to selection for other traits due to the close to zero genetic correlation between snout damage and the other recorded traits. However, before any selection against snout damage is performed the aetiology of the trait should be investigated to see if it can be reduced through management procedures including a better feed formulation.

The effect of the observed ulcer with increased mortality and skin wounds throughout the seawater phase may have masked differences between the three studied light regimes. However, to what extent this may have affected the reported results could not be determined.

## CRediT authorship contribution statement

**B. Gjerde**: Exp. design, statical analyses, writing the first draft, review and editing. **S.A. Boison**: Exp. design, review and editing. **D. Hazlerigg**: Conceptualization, funding acquisition, recording of data, review and editing. **T. Ytrestøyl**: Recording of data, review and editing. **T. Mørkøre**: Recording of data, review and editing. **E. Jørgensen**: Recording of data, review and editing. **A. Striberny**: Recording of data, review and editing. **S. R. Sandve**: Conceptualization, funding acquisition, project administration, review & editing.

## Data availability

Data will be made available on request.

